# BMP-3b suppresses proliferation, migration, invasion, and TGF-β 1/Smad3 signaling in breast cancer cells

**DOI:** 10.64898/2026.07.18.739378

**Authors:** Tatsuki Yaginuma, Kana Mizuta, Moyuri Inoue, Osamu Takahashi, Junpei Tanaka, Kazuya Haraguchi, Hiroki Tsurushima, Izumi Yoshioka, Shoichiro Kokabu, Manabu Habu

## Abstract

**Objective:** Bone morphogenetic protein-3b (BMP-3b), also known as growth differentiation factor 10, has been implicated in tumor suppression; however, its role in breast cancer and its interaction with transforming growth factor-β1 (TGF-β1) signaling remain incompletely understood.

**Methods:** Publicly available datasets were used to examine BMP-3b expression in breast lesions and its association with overall survival in patients with stage III or IV breast cancer. Human MCF-7 and murine 4T1 breast cancer cells were treated with recombinant BMP-3b. Cell proliferation, migration, invasion, epithelial–mesenchymal transition-related proteins, and TGF-β1-induced Smad3 phosphorylation were assessed using Cell Counting Kit-8, scratch wound-healing, Transwell invasion, and Western blot assays.

**Results:** BMP-3b expression was lower in ductal carcinoma in situ than in normal mammary tissue. Low BMP-3b expression was associated with poorer overall survival in patients with stage III or IV breast cancer. BMP-3b reduced proliferation of MCF-7 and 4T1 cells and inhibited migration and invasion of 4T1 cells. BMP-3b increased E-cadherin and decreased vimentin expression in both cell lines. It also attenuated TGF-β1-induced migration, invasion, and Smad3 phosphorylation in 4T1 cells.

**Conclusions:** BMP-3b suppresses malignant phenotypes of breast cancer cells and modulates TGF- β1/Smad3 signaling. These findings identify BMP-3b as a potential endogenous regulator of breast cancer progression.

## 1. Introduction

Breast cancer frequently metastasizes to bone, and skeletal metastases cause pain, pathological fractures, hypercalcemia, and other skeletal-related events that adversely affect quality of life and survival [1,2]. Bisphosphonates and anti-receptor activator of nuclear factor-κB ligand antibodies are widely used to prevent skeletal-related events and delay disease progression. However, these antiresorptive agents are associated with medication-related osteonecrosis of the jaw, which remains an important clinical problem in oral and maxillofacial practice [3,4]. A better understanding of breast cancer cell behavior and the factors that regulate malignant phenotypes may therefore be relevant to both oncological and oral healthcare.

The tumor microenvironment actively regulates cancer cell survival, proliferation, migration, and invasion. In bone, the extracellular matrix stores transforming growth factor-β (TGF-β), activins, and bone morphogenetic proteins (BMPs), which are released during bone remodeling [5–7]. TGF-β signaling is particularly important in breast cancer progression and can promote parathyroid hormone-related protein production, osteoclast activation, and tumor–bone interactions [8,9]. TGF-β also promotes epithelial–mesenchymal transition (EMT), migration, and invasion through canonical Smad2/3 signaling and non-canonical pathways.

BMP-3b, also known as growth differentiation factor 10, is a secreted member of the TGF-β superfamily with high sequence homology to BMP-3. BMP-3b is expressed in skeletal tissue and regulates cell differentiation and BMP signaling [10–12]. Reduced BMP-3b expression has been reported in several malignancies. In oral squamous cell carcinoma, loss of BMP-3b is associated with EMT, chemoresistance, and poor prognosis [13]. In triple-negative breast cancer cells, BMP-3b suppresses proliferation and EMT through upregulation of Smad7 [14]. Nevertheless, the clinical relevance of BMP-3b in breast cancer and its effects on TGF-β1-induced cellular responses require further clarification.

EMT is characterized by loss of epithelial markers such as E-cadherin and acquisition of mesenchymal markers such as vimentin and Snail, resulting in increased motility and invasiveness [15,16]. Previous work from our group demonstrated that TGF-β1/Smad3 signaling regulates EMT, migration, and invasion in oral squamous cell carcinoma cells [17]. We also showed that cytoskeletal regulatory proteins can influence tumor-cell proliferation, morphology, and adhesion [18]. These findings support the importance of examining both signaling and phenotypic changes when evaluating the effects of BMP-3b on cancer cells.

In the present study, we analyzed publicly available datasets to examine BMP-3b expression in breast lesions and its association with overall survival in patients with advanced breast cancer. We then investigated the effects of recombinant BMP-3b on proliferation, migration, invasion, EMT-related protein expression, and TGF-β1-induced Smad3 phosphorylation in human and murine breast cancer cells.

## 2. Materials and methods

### 2.1. Database analysis

Overall survival and BMP-3b expression data for patients with breast cancer were obtained from The Cancer Genome Atlas through the Human Protein Atlas. Patients with stage III or IV breast cancer were included. Patients were divided into high- and low-BMP-3b expression groups using the median expression value of 0.30 as the cutoff. Kaplan–Meier survival curves were generated, and differences between groups were evaluated using the log-rank test.

The Gene Expression Omnibus DataSet GDS3853, entitled “Ductal carcinoma in situ: mammary gland”, was used to analyze BMP-3b expression. The dataset was generated using the Affymetrix Human Genome U133 Plus 2.0 Array platform (GPL570). Expression was assessed using the GDF10 probe set 206159_at. Expression values and the associated P value were obtained from the GEO DataSet.

### 2.2. Cell culture

MCF-7 cells (RCB1904) were obtained from the RIKEN BioResource Research Center (Tsukuba, Japan). 4T1 cells (JCRB1447) were obtained from the Japanese Collection of Research Bioresources Cell Bank (Osaka, Japan). MCF-7 cells were cultured in Dulbecco’s modified Eagle’s medium, whereas 4T1 cells were cultured in RPMI 1640 medium. Both media were supplemented with 10% fetal bovine serum (Nichirei Biosciences, Tokyo, Japan), 100 units/mL penicillin, and 100 μg/mL streptomycin. Cells were maintained at 37°C in a humidified atmosphere containing 5% CO . For stimulation experiments, cells were cultured in medium containing 1% fetal bovine serum. Recombinant human BMP-3b and recombinant human TGF-β1 were used at final concentrations of 100 ng/mL and 5 ng/mL, respectively. Vehicle-treated cells were used as controls.

### 2.3. Reagents and antibodies

Recombinant human BMP-3b/GDF10 (Cat. No. 50165-M01H) was purchased from Sino Biological Inc. (Beijing, China). Recombinant human TGF-β1 was purchased from R&D Systems (Minneapolis, MN, USA). Cell Counting Kit-8 was purchased from Dojindo Laboratories (Kumamoto, Japan). Anti-phosphorylated Smad3 (Ser423/425; #9520) and anti-Smad2/3 (#8685) antibodies were purchased from Cell Signaling Technology (Danvers, MA, USA). Anti-E-cadherin antibody (#610181) was purchased from BD Biosciences (Franklin Lakes, NJ, USA). Anti-vimentin antibody (E-5; sc-373717) was purchased from Santa Cruz Biotechnology (Dallas, TX, USA). Anti-β-actin antibody (AC-15) was purchased from Sigma-Aldrich (St. Louis, MO, USA).

### 2.4. Cell proliferation assay

MCF-7 and 4T1 cells were seeded in 96-well plates at a density of 4 × 10³ cells/well. On the following day, cells were treated with BMP-3b or vehicle in medium containing 1% fetal bovine serum. MCF-7 cells were analyzed at 0, 24, and 48 h and at 7 and 14 days after treatment. 4T1 cells were analyzed at 0, 24, 48, and 72 h after treatment. Cell Counting Kit-8 reagent was added according to the manufacturer’s instructions, and absorbance at 450 nm was measured using a microplate reader (BD Biosciences, Franklin Lakes, NJ, USA).

### 2.5. Scratch wound-healing assay

The assay was performed with modifications to a previously described method [17]. 4T1 cells were seeded at a density of 5 × 10 cells/well in 24-well plates and cultured to confluence. A linear wound was created using a sterile pipette tip. Detached cells were removed by washing three times with phosphate-buffered saline. Cells were cultured in medium containing 1% fetal bovine serum and treated with vehicle, BMP-3b, TGF-β1, or BMP-3b plus TGF-β1. Phase-contrast images were obtained immediately after scratching and 24 h later. Wound areas were measured using ImageJ software (National Institutes of Health, Bethesda, MD, USA). The wound area at 0 h was defined as 100%.

### 2.6. Transwell invasion assay

The assay was performed with modifications to a previously described method [17]. 4T1 cells were suspended in serum-free medium and seeded in the upper chambers of Matrigel-coated Transwell inserts. Two hours after cell seeding, TGF-β1 was added to the lower chambers in the presence or absence of BMP-3b. The chambers were incubated for an additional 24 h. Cells remaining on the upper surface of the membranes were removed. Cells that had invaded to the lower surface were fixed and stained with crystal violet. Representative images were obtained using light microscopy.

### 2.7. Western blotting

Western blotting was performed as described previously [17,18]. Whole-cell lysates were separated by sodium dodecyl sulfate-polyacrylamide gel electrophoresis and transferred to polyvinylidene difluoride membranes. Membranes were incubated overnight at 4°C with primary antibodies diluted 1:1,000 in Tris-buffered saline containing Tween-20 (TBST) supplemented with 5% non-fat dry milk or bovine serum albumin and 0.01% sodium azide. Membranes were then incubated with horseradish peroxidase-conjugated secondary antibodies. Immunoreactive proteins were detected using enhanced chemiluminescence reagents (Cytiva, Marlborough, MA, USA).

For analysis of EMT-related proteins, MCF-7 and 4T1 cells were treated with BMP-3b or vehicle for 48 h. E-cadherin, vimentin, and β-actin were analyzed. For analysis of Smad3 phosphorylation, 4T1 cells were treated with TGF-β1 in the presence or absence of BMP-3b for 0, 10, 30, and 60 min. Phosphorylated Smad3, Smad2/3, and β-actin were analyzed.

### 2.8. Statistical analysis

Data are presented as the mean ± standard deviation from three independent experiments. Statistical analyses were performed using GraphPad Prism. Comparisons between two groups were performed using an unpaired Student’s t-test. Comparisons among multiple groups were performed using one-way analysis of variance followed by Fisher’s protected least significant difference post hoc test. Survival curves were compared using the log-rank test. P < 0.05 was considered statistically significant.

## 3.1. BMP-3b expression is reduced in breast cancer and is associated with prognosis in advanced breast cancer

Analysis of GEO DataSet GDS3853 showed that BMP-3b expression was significantly lower in ductal carcinoma in situ than in normal mammary tissue (Fig. 1A). Kaplan–Meier analysis of TCGA breast cancer data obtained through the Human Protein Atlas showed that, among patients with stage III or IV disease, the low-BMP-3b expression group had significantly poorer overall survival than the high-expression group (log-rank P = 0.019; Fig. 1B).

**Figure 1.**
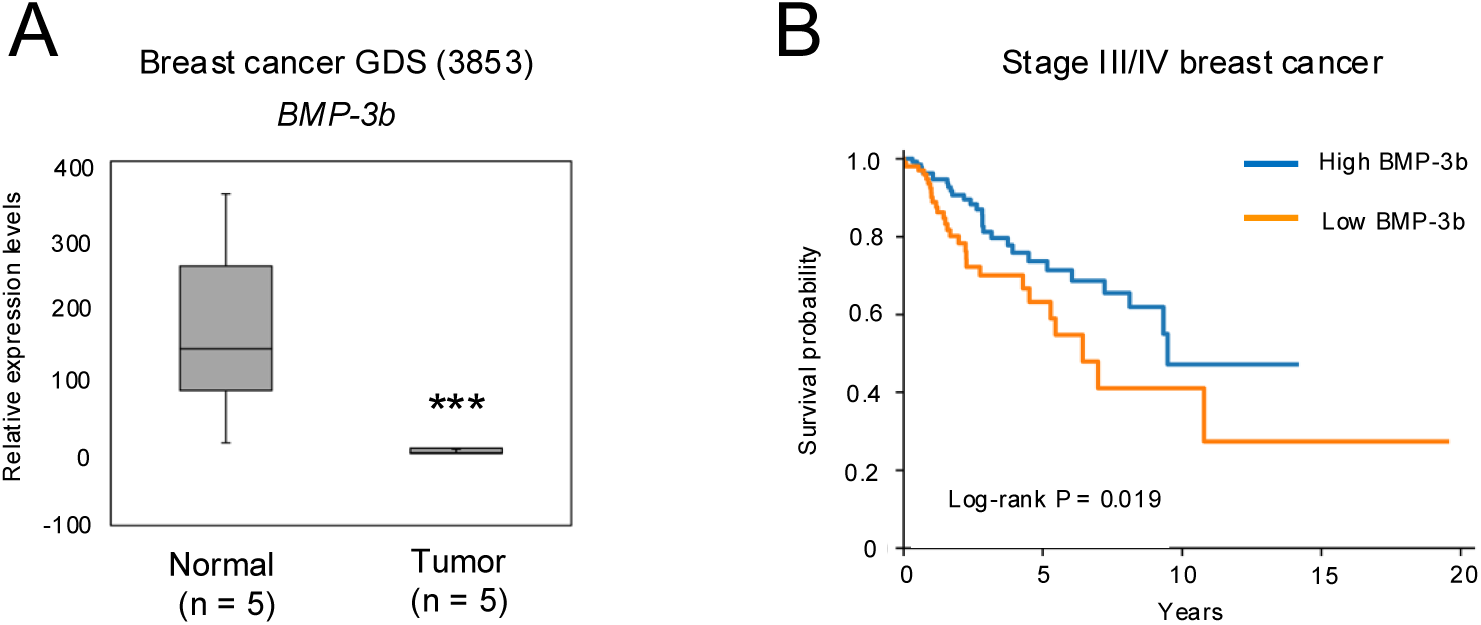
Reduced BMP-3b expression is associated with breast cancer and poor prognosis. (A) BMP-3b mRNA expression in normal mammary tissue and ductal carcinoma in situ was analyzed using Gene Expression Omnibus DataSet GDS3853. BMP-3b expression was significantly lower in ductal carcinoma in situ than in normal tissue. ***P < 0.001. (B) Kaplan–Meier analysis of overall survival in patients with stage III or IV breast cancer using The Cancer Genome Atlas data obtained through the Human Protein Atlas. Patients were stratified into high- and low-BMP-3b expression groups using the median expression value as the cutoff (0.30). Differences between groups were evaluated using the log-rank test (P = 0.019).

## 3.2. BMP-3b suppresses TGF-**β**1-induced migration, invasion, and Smad3 phosphorylation

BMP-3b reduced MCF-7 cell proliferation at 7 and 14 days, although this difference did not reach statistical significance, and significantly reduced 4T1 cell proliferation at 24 and 48 h compared with vehicle treatment (Fig. 2A,B). In the scratch wound-healing assay, BMP-3b significantly inhibited wound closure in 4T1 cells at 24 h (Fig. 2C,D). Western blot analysis showed that BMP-3b increased E-cadherin expression and decreased vimentin expression in both MCF-7 and 4T1 cells (Fig. 2E). TGF-β1 promoted wound closure in 4T1 cells, whereas co-treatment with BMP-3b significantly attenuated this response (Fig. 3A,B). In the Transwell invasion assay, fewer cells invaded through the Matrigel-coated membrane following combined treatment with TGF-β1 and BMP-3b than following TGF-β1 treatment alone (Fig. 3C). TGF-β1 induced Smad3 phosphorylation, whereas co-treatment with BMP-3b attenuated Smad3 phosphorylation without an evident change in total Smad2/3 protein levels (Fig. 3D).

**Figure 2.**
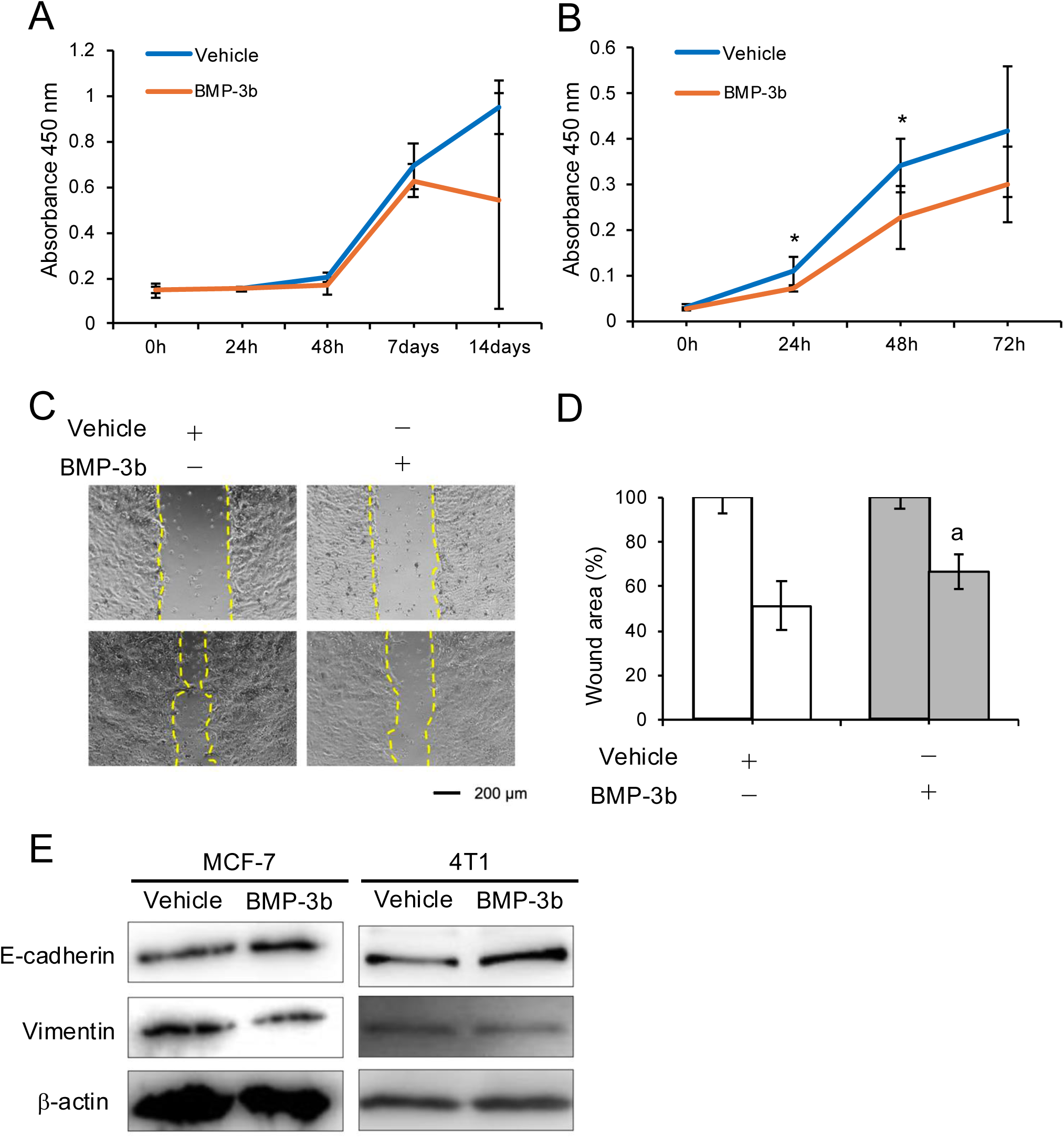
BMP-3b suppresses proliferation, migration, and epithelial–mesenchymal transition-related protein changes in breast cancer cells. (A,B) Proliferation of MCF-7 (A) and 4T1 (B) cells treated with vehicle or BMP-3b was evaluated using Cell Counting Kit-8. Absorbance at 450 nm was measured at the indicated time points. (C) Representative images of the scratch wound-healing assay in 4T1 cells treated with vehicle or BMP-3b (100 ng/mL). Images were obtained immediately after scratching and 24 h after treatment. Yellow dashed lines indicate the wound margins. Scale bar, 200 μm. (D) Quantification of the wound area shown in panel C. The wound area at 24 h was normalised to the initial wound area at 0 h. a, P < 0.05 versus the vehicle-treated group. (E) Western blot analysis of E-cadherin and vimentin in MCF-7 and 4T1 cells treated with vehicle or BMP-3b for 48 h. β-Actin was used as a loading control. Data are presented as the mean ± SD of three independent experiments. *P < 0.05 versus the vehicle-treated group at the corresponding time point.

**Figure 3.**
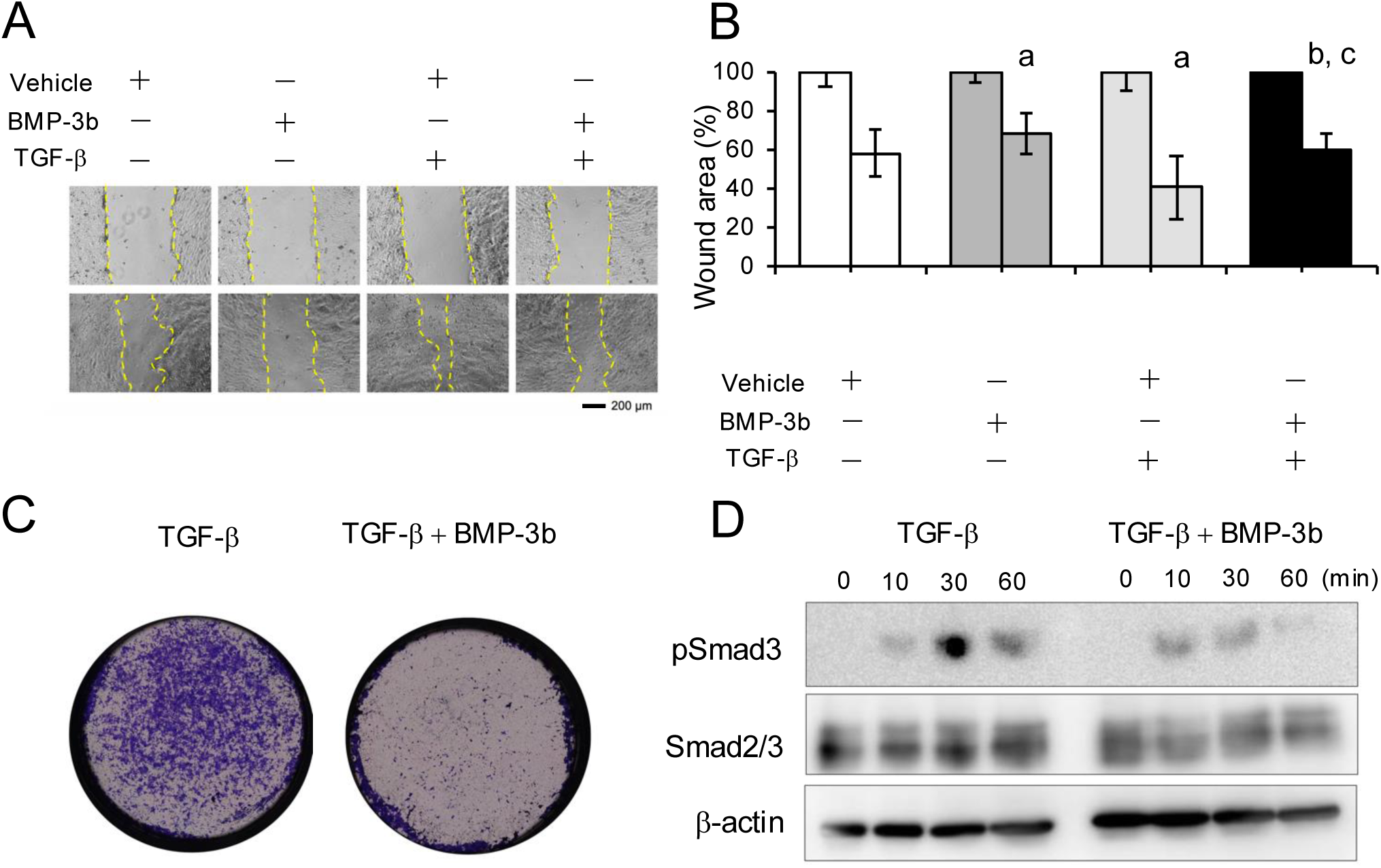
BMP-3b suppresses TGF-β1-induced migration, invasion, and Smad3 phosphorylation in 4T1 cells. (A) Representative images of the scratch wound-healing assay in 4T1 cells treated with vehicle, BMP- 3b (100 ng/mL), TGF-β1 (5 ng/mL), or BMP-3b plus TGF-β1. Images were obtained immediately after scratching and 24 h after treatment. Yellow dashed lines indicate the wound margins. Scale bar, 200 μm. (B) Quantification of the wound area shown in panel A. The wound area at 24 h was normalised to the initial wound area at 0 h. a, P < 0.05 versus the vehicle-treated group; b, P < 0.05 versus the BMP-3b-treated group; c, P < 0.05 versus the TGF-β1-treated group. (C) Representative images of the Transwell invasion assay in 4T1 cells treated with TGF-β1 alone or with TGF-β1 plus BMP-3b. Invaded cells were stained with crystal violet. (D) Western blot analysis of phosphorylated Smad3 and total Smad2/3 in 4T1 cells treated with TGF-β1 (5 ng/mL) alone or in combination with BMP-3b (100 ng/mL) for 0, 10, 30, and 60 min. β-Actin was used as a loading control. Quantitative data are presented as the mean ± SD of three independent experiments.

## 4. Discussion

The present study showed that BMP-3b expression was reduced in breast cancer tissue and that low BMP-3b expression was associated with poorer overall survival in patients with stage III or IV breast cancer. Recombinant BMP-3b suppressed proliferation of human and murine breast cancer cells and inhibited migration and invasion of 4T1 cells. BMP-3b also increased E-cadherin expression, decreased vimentin expression, and attenuated TGF-β1-induced Smad3 phosphorylation. These findings support a role for BMP-3b as a negative regulator of malignant phenotypes in breast cancer cells.

BMP-3b is a secreted TGF-β superfamily member involved in the regulation of cell differentiation and BMP signaling [10–12,19]. Reduced BMP-3b expression has been reported in several malignancies. Cheng et al. showed that loss of BMP-3b was associated with EMT, chemoresistance, and poor prognosis in oral squamous cell carcinoma [13], while Zhou et al. demonstrated that BMP-3b suppressed proliferation and EMT in triple-negative breast cancer cells through Smad7 upregulation [14]. The present database analyses and cell-based findings are consistent with these reports and further support the tumor-suppressive properties of BMP-3b in breast cancer.

EMT contributes to cancer-cell migration and invasion and is characterized by decreased E-cadherin and increased vimentin expression [15,16]. In the present study, BMP-3b increased E-cadherin, decreased vimentin, and suppressed migration and invasion. These effects indicate that BMP-3b regulates both proliferative and motile phenotypes of breast cancer cells. Our previous studies showed that TGF-β1/Smad3 signaling promotes EMT, migration, and invasion in oral squamous cell carcinoma cells [17] and that cytoskeletal organization is closely linked to tumor-cell proliferation, morphology, and adhesion [18]. Together, these observations emphasize that changes in EMT-related proteins and cell motility are integral components of the BMP-3b response.

TGF-β is a major inducer of EMT and promotes migration and invasion through Smad2/3-dependent signaling [15–17]. Here, BMP-3b attenuated TGF-β1-induced migration, invasion, and Smad3 phosphorylation. BMP-3b has been reported to activate Smad2/3 signaling in some cell types [11,19], whereas recent evidence indicates that it can also antagonize BMP receptor signaling. Kodama et al. demonstrated that BMP-3b interferes with the interaction between BMP4 and BMP type I receptors and suppresses BMP4-induced Smad1/5 phosphorylation [12]. These findings suggest that BMP-3b does not simply activate a single pathway but may modulate TGF-β superfamily signaling according to cell type, receptor expression, and the presence of other ligands. The mechanism by which BMP-3b attenuates TGF-β1-induced Smad3 phosphorylation in breast cancer cells remains to be determined.

This study has several limitations. First, the experiments were primarily performed in two breast cancer cell lines, and the effects of BMP-3b in other molecular subtypes remain unknown. Second, the effects of BMP-3b on tumor growth and invasion were not examined in vivo. Third, only E-cadherin and vimentin were examined as EMT-related proteins. Finally, the receptor-level mechanism underlying the inhibition of Smad3 phosphorylation was not investigated. Further studies using additional breast cancer models and receptor-interaction analyses are required.

## 5. Conclusions

BMP-3b expression was reduced in breast cancer tissue, and low expression was associated with poorer overall survival in patients with advanced breast cancer. BMP-3b suppressed breast cancer cell proliferation, migration, and invasion and modulated E-cadherin, vimentin, and TGF-β1-induced Smad3 phosphorylation. BMP-3b may therefore function as an endogenous regulator of malignant phenotypes in breast cancer cells.

## Data availability

The data that support the findings of this study are available from the corresponding author upon reasonable request. Publicly available data were obtained from the Gene Expression Omnibus, The Cancer Genome Atlas, and the Human Protein Atlas.

## CRediT authorship contribution statement

**T.Y.: Conceptualization, methodology, investigation, formal analysis, writing—original draft, and funding acquisition.**

K.M.: Methodology, investigation, and data curation.

O.T.: Investigation and data curation. J.T.: Investigation and data curation. K.H.: Investigation and data curation. H.T.: Investigation and data curation.

M.I.: Formal analysis, visualization, and writing—review and editing. I.Y.: Supervision and writing—review and editing.

S.K.: Conceptualization, supervision, project administration, funding acquisition, and writing— original draft, review, and editing.

M.H.: Supervision and writing—review and editing.

All authors reviewed and approved the final manuscript.

## Declaration of generative AI and AI-assisted technologies in the writing process

During the preparation of this work, the authors used ChatGPT (OpenAI) to assist with English-language editing and manuscript organization. The authors subsequently reviewed and edited the content and take full responsibility for the content of the publication.

